# MicroRNA childhood Cancer Catalog (M3Cs): A Resource for Translational Bioinformatics Toward Health Informatics in Pediatric Cancer

**DOI:** 10.1101/2021.07.26.452794

**Authors:** Wafaa M. Rashed, Fatima Adel, Mohamed A. Rezk, Lina Basiouny, Ahmed A. Rezk, Ahmed H. Abdel-Razek

## Abstract

**Background:** MicroRNA childhood Cancer Catalog (M3Cs) is high-quality curated collection of published miRNA research studies on 16 pediatric cancer diseases. M3Cs scope was based on two approaches: data-driven clinical significance and data-driven human pediatric cell line models.

**Method:** M3Cs development passed through three phases: 1. Literature Mining: It includes external database search and screening. **2**. Data processing that includes 3 steps: a-Data Extraction, b-Data Curation & annotation, c-Web Development. **3**. Publishing: Shinyapps.io was used as a web interface for the deployment of M3Cs. M3Cs is now available online and can be accessed through https://m3cs.shinyapps.io/M3Cs/.

**Results:** For Data-driven clinical significance Approach, 538 miRNAs from 268 publications were reported in the clinical domain while 7 miRNAs from 5 publications were reported in the clinical & drug domain. For data-driven human pediatric cell line models approach, 538 miRNAs from 1268 publications were reported in cell line domain while 211 miRNAs from 177 publications in cell line & drug domain.

**Conclusion:** M3Cs acted to fill the gap by applying translational bioinformatics (TBI) general pathway to transfer data-driven research toward data-driven clinical care and/or hypothesis generation. Aggregated and well-curated data of M3Cs will enable stakeholders in health care to incorporate miRNA in the clinical policy.

## Introduction

Although pediatric cancer diseases are rare, they represent a significant cause of both mortality and morbidity (Johnston et al., 2021). Favorable outcomes of pediatric cancer diseases require effective risk assessment and early diagnosis followed by personalized treatment. By harnessing the power of early diagnosis and personalized treatments in pediatric cancer diseases, the overall healthcare system outcome will be improved, and the cost of therapy will be lowered.

MicroRNAs (miRNAs) are small RNA molecules (≈ 22 nucleotides) that belong to the class of non-coding RNA (Rashed et al., 2019). They play an important role in post-transcriptional regulation of gene expression via mRNA degradation and/or translational repression (Catalanotto et al., 2016). Research has accumulated a large body of evidence implicating many miRNAs in the diagnosis, prognosis and risk of hematological and solid tumors of childhood cancer (Carvalho de Oliveira et al., 2018; Gulino et al., 2015; Leichter et al., 2017; Murray et al., 2015; Smith et al., 2019). Using bioinformatics, there are about seven categories of online miRNA databases, however none of which includes specialized miRNA databases for pediatric cancers (Shaker et al., 2020).

Great advances in biomedical research (e.g., high throughput sequencing and functional analysis) have led to voluminous data that needs proper processing into information then transformation into knowledge applied in the clinical setting. This will fill the gap between bench and bedside and enable clinicians to achieve favorable outcomes of pediatric cancer diseases. Translational Bioinformatics (TBI) is one of the biomedical research channels that can play an inevitable role in this aspect. Through convergence of molecular bioinformatics, clinical informatics, genetics, biostatics and health informatics, recently many TBI databases have been established (Singh et al., 2019). Not only did these databases create new knowledge, but also, they provided novel insights into the disease genetic mechanism. Database for miRNA of pediatric cancer will structure the microRNA clinical utility knowledge in pediatric cancer and consequently, it may help in utilizing miRNA research findings into the clinical side.

That is why we have launched MicroRNA Childhood Cancer Catalog (M3Cs). Based on the translational bioinformatics (TBI) spectrum, the principle of this platform is to bring miRNA research into clinical significance in both pediatric cancer patient care and drug discovery and toward health informatics in childhood cancer. This will ease the policy makers’ exploitation of pediatric cancer patients’ data to shift emphasis to prevention, allow the selection of optimal diagnosis and therapy and reduce trial and error prescribing as well as reviving drugs that failed early in clinical trials. Additionally, it will help scientists for more hypothesis generation. To achieve this goal, we have utilized two approaches; data-driven clinical significance and data-driven human pediatric cell line models. (Figure 1)

**Figure 1:**
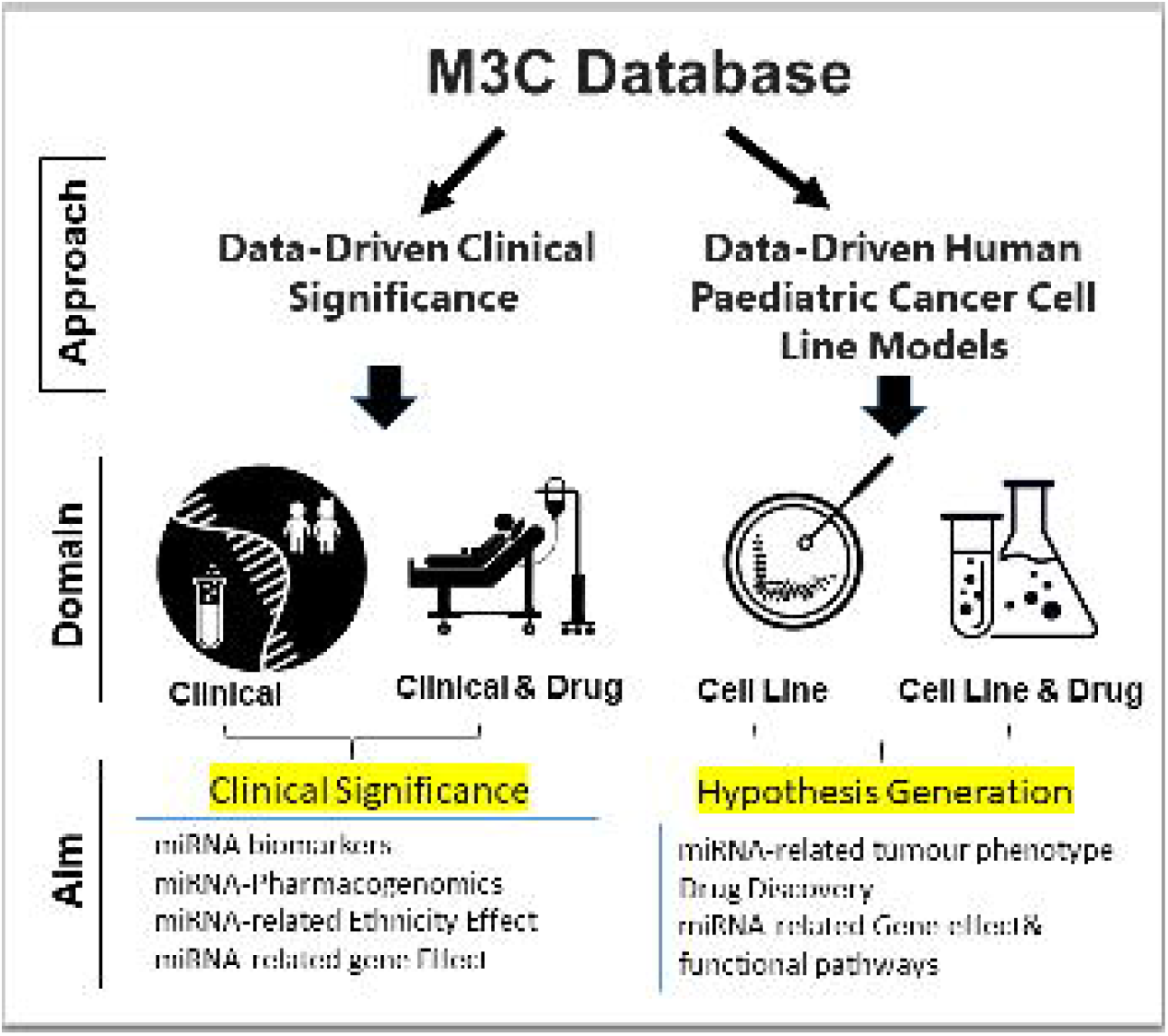
M3Cs approaches, domains and aims.

## Method

M3Cs development passed with three phases with central validation at the end of each phase. (**Figure 2**)

**Figure 2:**
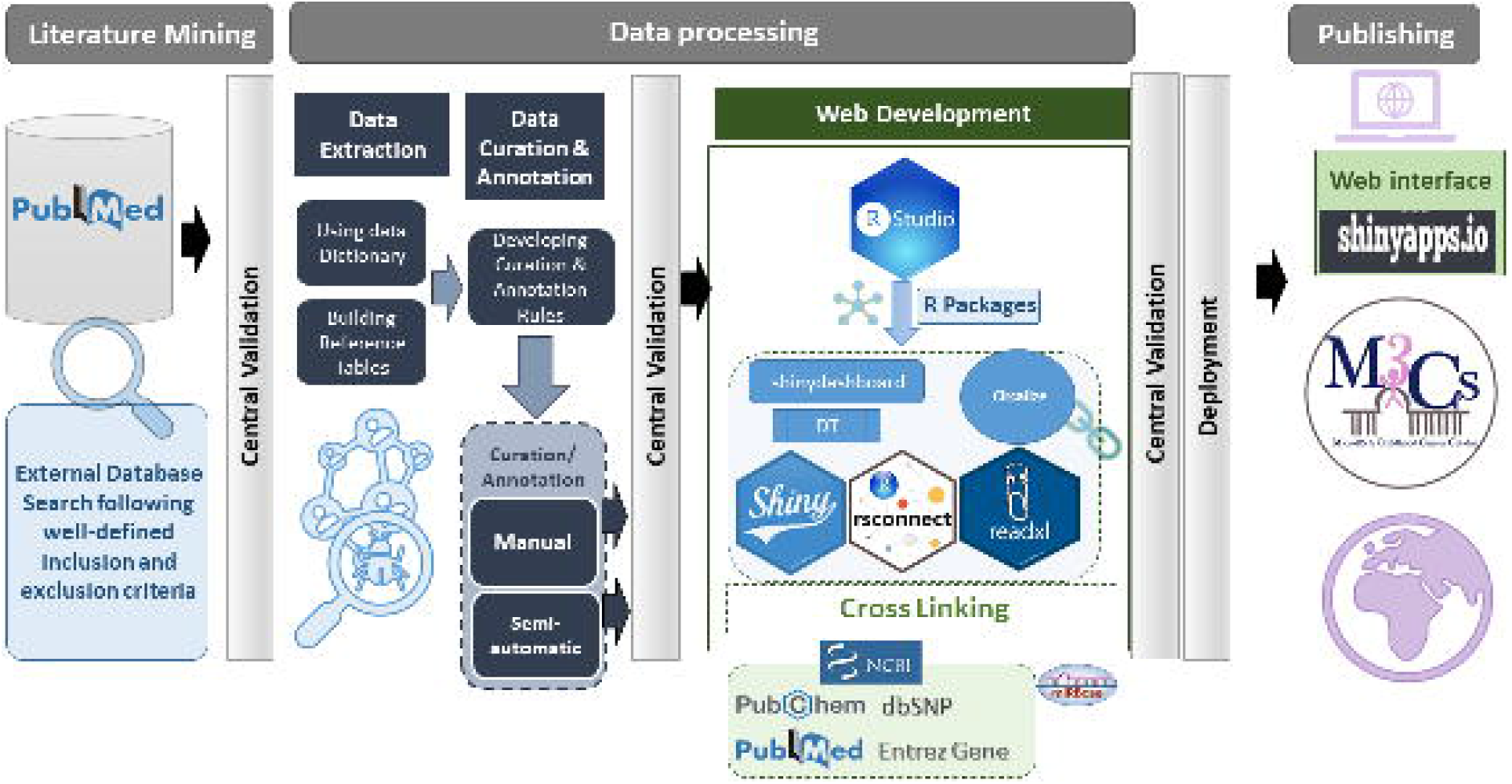
The Overall Workflow of M3Cs phases.

### Phase 1: Literature Mining: It included external database search and screening

M3Cs team was responsible for searching PubMed using the following search terms: (((microRNA) OR miRNA) OR miR) AND “Disease name”). We have included the most common pediatric cancer diseases: Acute Lymphoblastic Leukemia (ALL), Acute Myeloid Leukemia (AML), Adrenocortical Tumor (ACT), Brain Tumor, Chronic Myelogenous leukemia (CML), Ewing Sarcoma (ES), Hepatoblastoma, Hodgkin disease, Juvenile myelomonocytic leukemia (JMML), Langerhans cell histiocytosis (LCH), Neuroblastoma (NB), Non-Hodgkin lymphoma (NHL), Osteosarcoma (OS), Renal tumor, Retinoblastoma (RB), Rhabdomyosarcoma (RMS). Screening of search results was based on inclusion and exclusion criteria.

#### For data driven clinical significance approach

the inclusion criteria for studies are: (1) Patients with age less than 18 years; (2) Publication with data about miRNA related-disease diagnosis, -disease prognosis, or -disease risk. For publications of prognostic miRNAs, survival results were measured, and it should include at least one survival curve [overall survival (OS), disease-free survival (DFS), recurrence/relapse-free survival (RFS), progression-free survival (PFS), and metastasis-free survival (MFS)] with or without HRs/95% CIs. MiRNA studies were excluded if: (1). Age of patients was not identified or above 18 years old; (2). Reports or reviews or letters without primary data; (3). Non-English publication; (4). Retracted papers.

#### For data-driven human pediatric cell line models approach, the inclusion criteria of miRNA studies are (1)

Human cell line of pediatric origin (<18 years) with verification from Cellosaurus(Bairoch, 2018) [species origin and age at sampling]; (2). Publication investigated the miRNA Effect on the tumor phenotype. MiRNA studies were excluded if: (1). Data from animal models; (2). Reports, reviews, or letters without primary data; (3). Non-English publications; (4). Retracted papers. Central Validation is the safeguard for M3Cs. Centralization of data validation decreased variability and ensured database of high quality. The primary role of central validation at the end of phase 1 was to revise all included and excluded studies using the predefined criteria. Disputes regarding including or excluding studies have been resolved by discussion with the whole team.

### Phase 2: Data processing that included 3 steps: 1-Data Extraction, 2-Data Curation & annotation, 3-Web Development

#### Step 1-Data Extraction

M3Cs is based on two main approaches: data driven clinical significance approach and data-driven human pediatric cell line model approach. Each approach has two specific domains. For data driven clinical significance approach, there is “clinical domain” and “clinical & drug domain”. For data-driven human pediatric cell line model approach, there is “cell line domain” and “cell line & drug domain”. M3Cs established data collection forms for each domain containing a recommended set of variables of data elements. These data collection forms are used to collect research data from each eligible study for the M3Cs database. Data dictionary for each domain was developed with the standardized data definitions and formats for the collection. (Supplementary tables 1,2,3 & 4).

#### Step 2-Data Curation & Annotation

Data curation was done manually by well-trained scientists. Many online databases were used for reporting of some variables [e.g., miRBASE (Kozomara et al., 2019), Entrez gene (Brown et al., 2015), PubChem (Kim et al., 2020), dbSNP (Sherry et al., 2001) and PubMed (Fiorini et al., 2018)]. Also, in the data driven clinical significance approach, we followed GWAS catalog standard method of reporting ancestry data (Morales et al., 2018). At the end of this phase, **central validation** was used to check both the quality and accuracy of manually curated and annotated data.

**Table 1.** Summary of the total number of included and excluded studies for each disease in the overall phases of M3Cs.

| Serial | Disease | Included | Excluded | Total Screened |
| --- | --- | --- | --- | --- |
| 1 | Acute Lymphoblastic Leukemia | 108 | 227 | **335** |
| 2 | Acute Myeloid Leukemia | 115 | 911 | 1026 |
| 3 | Adrenocortical Tumors | 1 | 73 | 74 |
| 4 | Brain Tumors | 78 | 233 | **311** |
| 5 | Chronic Myelogenous Leukemia | 4 | 314 | **318** |
| 6 | Ewing Sarcoma | 22 | 50 | **72** |
| 7 | Hepatoblastoma | 149 | 51 | **200** |
| 8 | Hodgkin Disease | 1 | 226 | **227** |
| 9 | Juvenile Myelomonocytic leukemia (JMML) | 3 | 5 | **8** |
| 10 | Langerhans cell Histiocytosis (LCH) | 0 | 0 | **0** |
| 11 | Neuroblastoma | 160 | 416 | **576** |
| 12 | Non-Hodgkin Lymphoma (NHL) | 38 | 622 | **660** |
| 13 | Osteosarcoma | 1084 | 270 | 1354 |
| 14 | Rhabdomyosarcoma | 31 | 102 | **133** |
|  | (RMS) |  |  |  |
| 15 | Renal Tumor | 14 | 1131 | **1145** |
| 16 | Retinoblastoma | 64 | 39 | **103** |
| Total |  | 1872 | 4670 | **6542** |

#### Step 3-Web Development

M3Cs is a web application that has been developed by R programming language (version 4.0.2) and RStudio was used as an interface (https://www.r-project.org) and Shiny framework was used for deployment of the application (https://CRAN.R-project.org/package=shiny). The data of each domain were presented using the DT package, while the miRNA plots were depicted with the circlize package. We have used crosslink to ease access to other websites. The rsconnect package was used for deployment of the shiny application. At the end of this phase, **central validation** was responsible for checking high quality cross links.

### Phase 3: Publishing

Shinyapps.io platform was used as a web interface for deployment of M3Cs application. Shinyapps.io platform is secure by design and allows M3Cs shiny application to run in its own protected environment.

## Result

M3Cs can be searched using the Web interface https://m3cs.shinyapps.io/M3Cs/. The homepage of M3Cs includes a disclaimer that gives important notices for users related to many issues: the content, the medical information and advice, the endorsement, the intellectual property, as well as the external links. Out of 6542 studies screened from PubMed between 2002 till the end of 2020, 1872 studies were included in the M3Cs. The total number of included and excluded studies in each disease in M3Cs were summarized in table 1. In phase 1 (literature mining), we did not find any miRNA related LCH study in pediatric cancer patients.

The M3Cs manual page includes a data dictionary for each M3Cs domain (clinical, clinical & Drug, Cell line, Cell line & Drug) to give the definitions of different variables. The main M3Cs search page includes the four domains. M3Cs allows retrieving information of candidate miRNA in each domain. Also, it can be searched with different possible searches due to using filter tables in each domain. **(Figure 3)**

**Figure 3.**
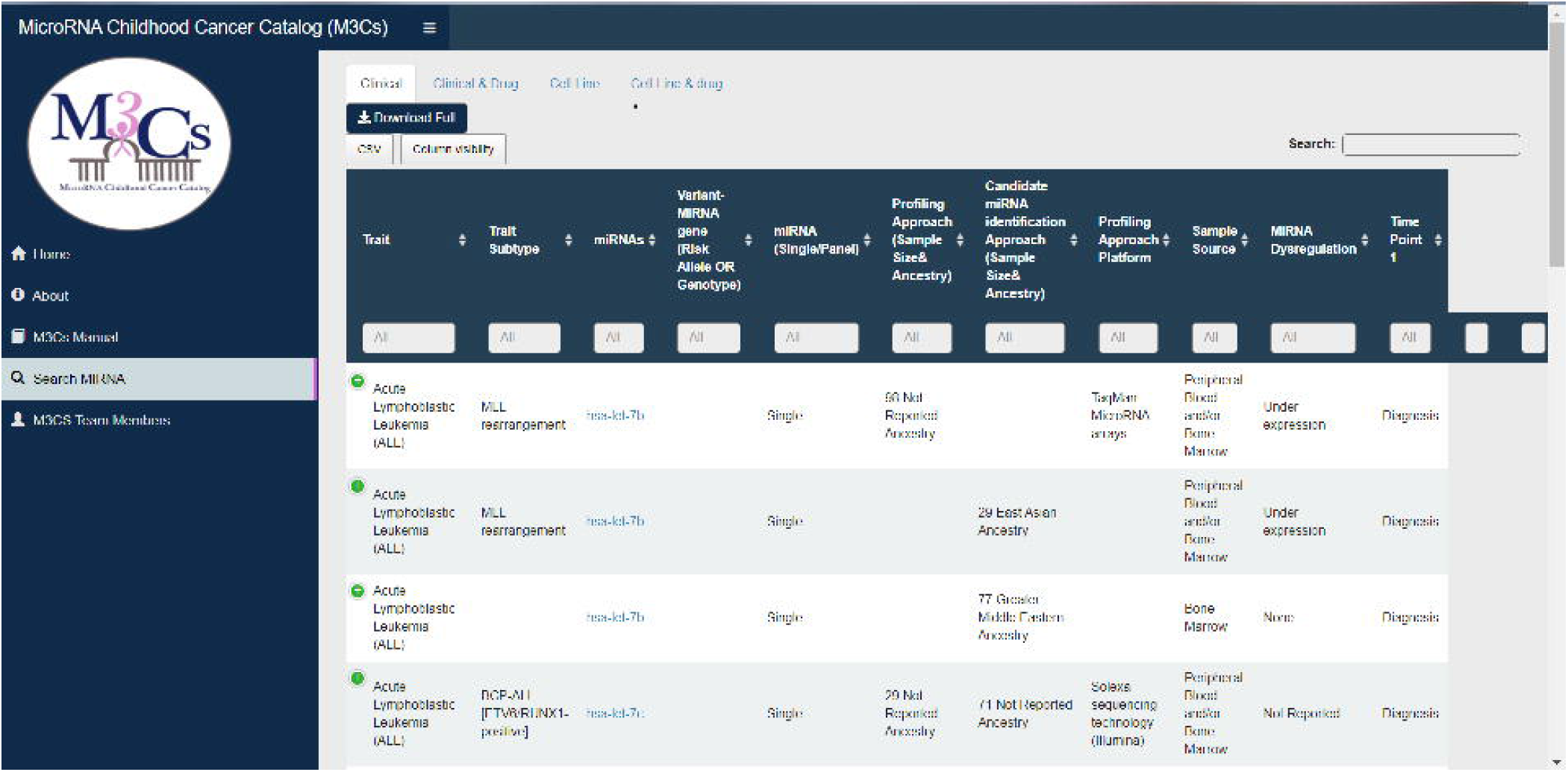
Screenshot of search miRNA page in M3Cs.

### I- Data-Driven Clinical Significance Approach

In the clinical domain, miRNA role as a probable diagnostic, prognostic, predictive biomarker along with miRNA-Polymorphism value as a probable risk factor for pediatric cancer trait and trait subtype has been reported.

M3Cs helps to identify reported miRNAs that can be used as a biomarker panel (a group of miRNAs or miRNAs & genes). M3Cs determines the study design approach (profiling approach then candidate approach (as a validation) or profiling approach only or candidate approach only). In the clinical & drug domain, M3Cs reports clinical studies related to miRNAs-associated drug toxicity in pediatric cancer traits and trait subtypes. Variants in miRNA gene and drug toxicity associated with this have been reported. In both domains, sample size and ethnicity were described. Pediatric patients’ ethnicity was reported following the GWAS catalog framework(Morales et al., 2018). In addition, miRNA dysregulation and their tested direct downstream target gene(s) have been reported. All statistical significant parameters described by the original study have been included by M3Cs e.g. Hazard ratio, covariates, odds Ratio, relative risk, sensitivity and specificity.

#### Sample Data from retinoblastoma studies

##### Data description of clinically diagnostic miRNA biomarkers in Retinoblastoma

Supplementary table 5 and Supplementary figure 1 represent data of clinically diagnostic miRNA biomarkers in 22 retinoblastoma studies. One study recruited subjects from American population, while the remaining studies recruited subjects of Asian population. All these diagnostic miRNAs have been reported from retinoblastoma tissues except three miRNAs (hsa-let-7e-5p, hsa-miR-21-5p and hsa-miR-320a-3p) that reported from peripheral blood. Out of the 40 miRNAs, 22 miRNAs showed overexpression while 18 miRNAs showed under-expression.

Also, 31 miRNAs can be used as a single diagnostic biomarker while 8 miRNAs can be used as a panel biomarker. Five miRNAs, known as miR-17-92 cluster, can be used as diagnostic biomarker panel while each of the remaining three miRNAs (hsa-let-7e-5p, hsa-miR-21-5p and hsa-miR-320a-3p) can be used as a panel in combination of each one with *NSE* gene.

##### Pathway Enrichment Analysis and Intersected Target Genes

We used DIANA-TarBase v7.0 of DIANA-miRPath v3.0 (Vlachos et al., 2015) for better understanding of the functional role of the clinically diagnostic miRNAs identified in retinoblastoma studies. We entered all clinically diagnostic miRNAs in the search bar of DIANA-miRPath v3.0 to identify intersected genes in the Kyoto Encyclopedia of Genes and Genomes (KEGG) pathways and Gene Ontology (GO) categories. To identify the significant KEGG pathways modulated by the same miRNAs, we created HeatMap of diagnostic miRNAs vs pathways (supplementary figure 2).

Using intersected genes modulated by the 40 diagnostic miRNAs, only two significant associated KEGG pathways were identified by DIANA-TarBase v7.0 of DIANA-miRPath V3.0. both *BTG2* and *DICER1* are the intersected gene in the RNA degradation (hsa03018) & microRNAs in cancer (hsa05206) in KEGG pathways, respectively (supplementary table 6). Supplementary table 7 represents the significant GO molecular function and the intersected genes that are modulated by the selected clinically diagnostic miRNAs as identified by DIANA-TarBase v7.0 of DIANA-miRPath V3.0. Using the 40 diagnostic miRNAs biomarker in retinoblastoma, the same interested genes in KEGG pathways (*BTG2* & *DICER1*) have been identified in GO molecular function in addition to *AHNAK* gene.

### II - Data-Driven Human Pediatric Cancer Cell Line Model Approach

In the cell line domain, M3Cs included most human pediatric cell line studies that identify the effect of dysregulated miRNA on the tumor phenotype in each pediatric cancer trait and trait subtype to differentiate between miRNAs that act as a tumor suppressor or Oncogene. In cell Line & Drug Domain, studies examining the drug Effect on miRNA expression in each pediatric cancer trait and trait subtype have been included. In both domains direct downstream target genes associated with this miRNA along with the underlying affected downstream pathways have been reported. Publication title, PubMed and Notes were the three fixed variables included in all domains.

If M3Cs team missed specific publications about microRNA in pediatric cancer, users are welcome to send us using M3Cs contact email (**)** and the M3Cs team is pleased to add them to the M3Cs. Using the same email, M3Cs users can contact M3Cs to draw circos plots to visualize miRNA.

## Discussion

M3Cs is the first specialized miRNA pediatric cancer database. M3Cs utilized two approaches; data-driven clinical significance and data-driven human pediatric cell line model approaches. In data-driven clinical significance, M3Cs assembled studies that investigate the potential role of miRNA as a diagnostic, prognostic and predictive biomarker in each pediatric cancer disease. In addition, it promotes researchers to use data to investigate a new biomarker panel and to pursue more meta-analysis studies. Furthermore, it can be used as a springboard for drug discovery and gene therapy. Moreover, it reports miRNA-Polymorphism that affects drug response (miRNA Pharmacogenomics). This will pay attention to miRNA pharmacogenomics to be used in contemporary treatment protocols and consequently more research in this promising field.

Furthermore, M3Cs reports clinical studies with dysregulated miRNAs and their tested direct target gene(s). It enables researchers to fuse diagnostics with targeted therapy as personalized medicines to improve both the management and outcome of many pediatric cancer diseases.

In data-driven human pediatric cell line model approach, M3Cs shed light on miRNA value as a targeted therapy. Developing strategies to replenish tumor suppressive miRNAs (miRNA replacement) or supprese oncomiRs (anti-miRNAs) can be utilized as therapeutic modulation of miRNAs. Besides, developing chemical modification & delivery systems for these miRNA-based therapeutics have been introduced *in vivo* cancer models (Rupaimoole & Slack, 2017). All these factors will encourage researchers to develop *in vivo* models to treat different types of pediatric cancers.

Also, M3Cs will help computer scientists to use the curated data in creating simulation for easy biomarker discovery and modeling of any biological system using big data.

M3Cs have developed variables of datasets that can be used by the journal editors and reviewers to evaluate submitted miRNA articles. We have excluded many publications that include missing parts in analysis that can be included upon reviewers’ request. Standardization of reporting in miRNA publications will help researchers to advance miRNA research and applications in terms of increasing the number of evidence.

There will be a half-annual update of M3Cs that will include adding new publications and removing any retracted paper from M3Cs records.

### Future plan

M3Cs survey has been developed to help users to send their feedback about M3Cs design, content and ease of search. M3Cs version 2 (M3Cs v.2) will take in consideration M3Cs users’ feedback as well as the inclusion of more pediatric cancer types and other features.

In M3Cs, we followed the general pathway of translational bioinformatics (TBI) of data collection, storage, analysis, retrieval, and interpretation. We acted to fill the gap by applying TBI which simply transfers data-driven research toward data-driven clinical care and/or hypothesis generation. Information that is aggregated and well-curated will enable stakeholders in health care to incorporate miRNA in the clinical policy to be used by clinicians in their clinics.

## Supplementary Tables

**Supplementary Table 1**. Clinical Domain and Associated Variables.

**Supplementary Table 2**. Clinical & Drug Domain and Associated Variables.

**Supplementary Table 3**. Cell Line Domain and Associated Variables.

**Supplementary Table 4**. Cell Line & Drug Domain and Associated Variables

**Supplementary Table 5**. Forty miRNAs as probable clinically diagnostic biomarkers in Retinoblastoma in M3Cs.

**Supplementary Table 6**. Summary of the statistically significant molecular KEGG pathways and intersected genes that are modulated by the selected clinically diagnostic miRNAs in retinoblastoma patients.

**Supplementary Table 7**. Summary of the significant GO molecular function and intersected genes that are modulated by the selected clinically diagnostic miRNAs in retinoblastoma patients.

## Supplementary Figures

**Supplementary Figure 1**. Circos plot of probable diagnostic miRNA in Retinoblastoma.

**Supplementary Figure 2**. Heatmap of KEGG pathways vs clinically diagnostic miRNAs in retinoblastoma.

## Authors’ Contribution

**WMR:** Conceptualization, Conception and design and writing original draft. MAR, FA, LB, AAR and AHA: literature mining, data extraction and data curation. **ASA**: Web development. **WMR, MAR, FA, LB, AAR and AHA**: Writing-Reviewing and Editing, final approval of article.

## Acknowledgements

M3Cs team would like to thank Professor Dr. Logan Spector, University of Minnesota-US, for his valuable scientific consultation and appreciated support along with the phases of developing this platform. Also, M3Cs team would like to thank Mohamed A. Hussein, Ahmed Ismail, and Nancy Sherif for their participation in phase 1 of this catalog.

