## Supplementary figures and images for "MicroRNA childhood Cancer Catalog (M3Cs): A Resource for Translational Bioinformatics Toward Health Informatics in Pediatric Cancer"

### Supplementary Figure 1

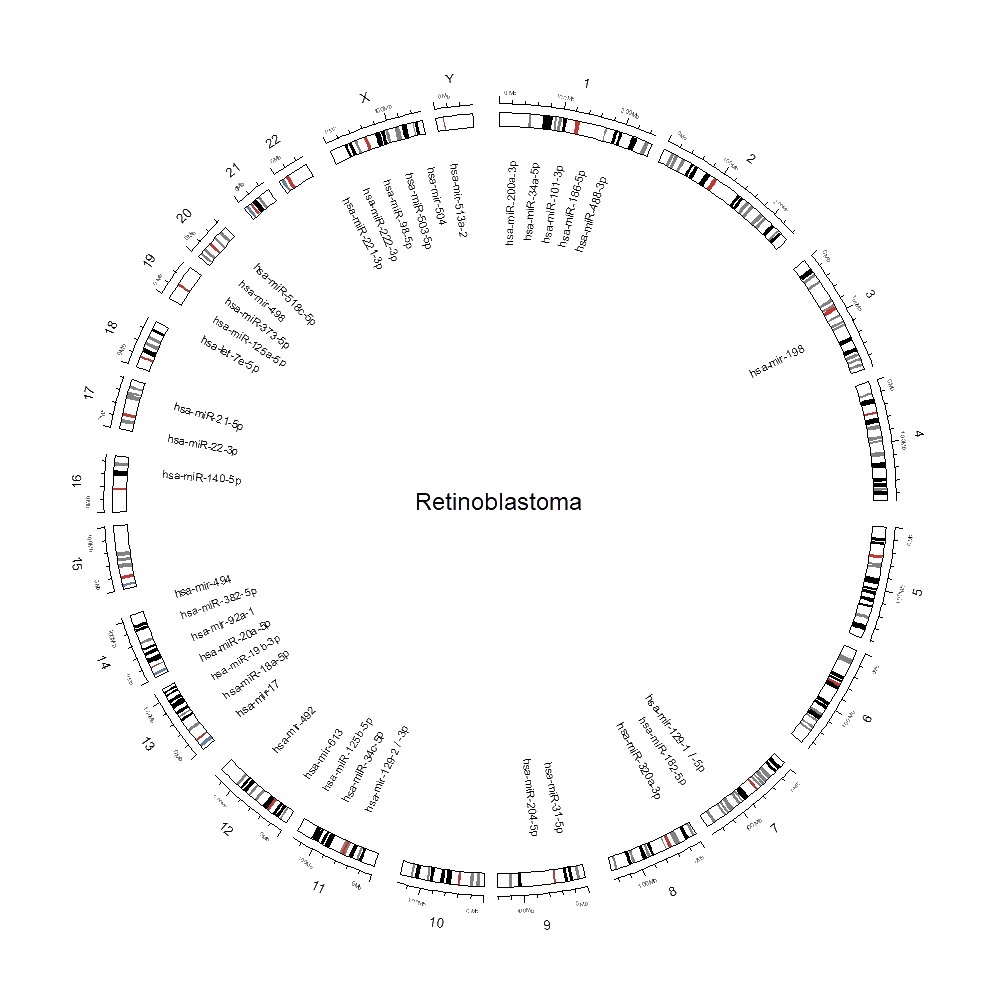

### Supplementary Figure 2

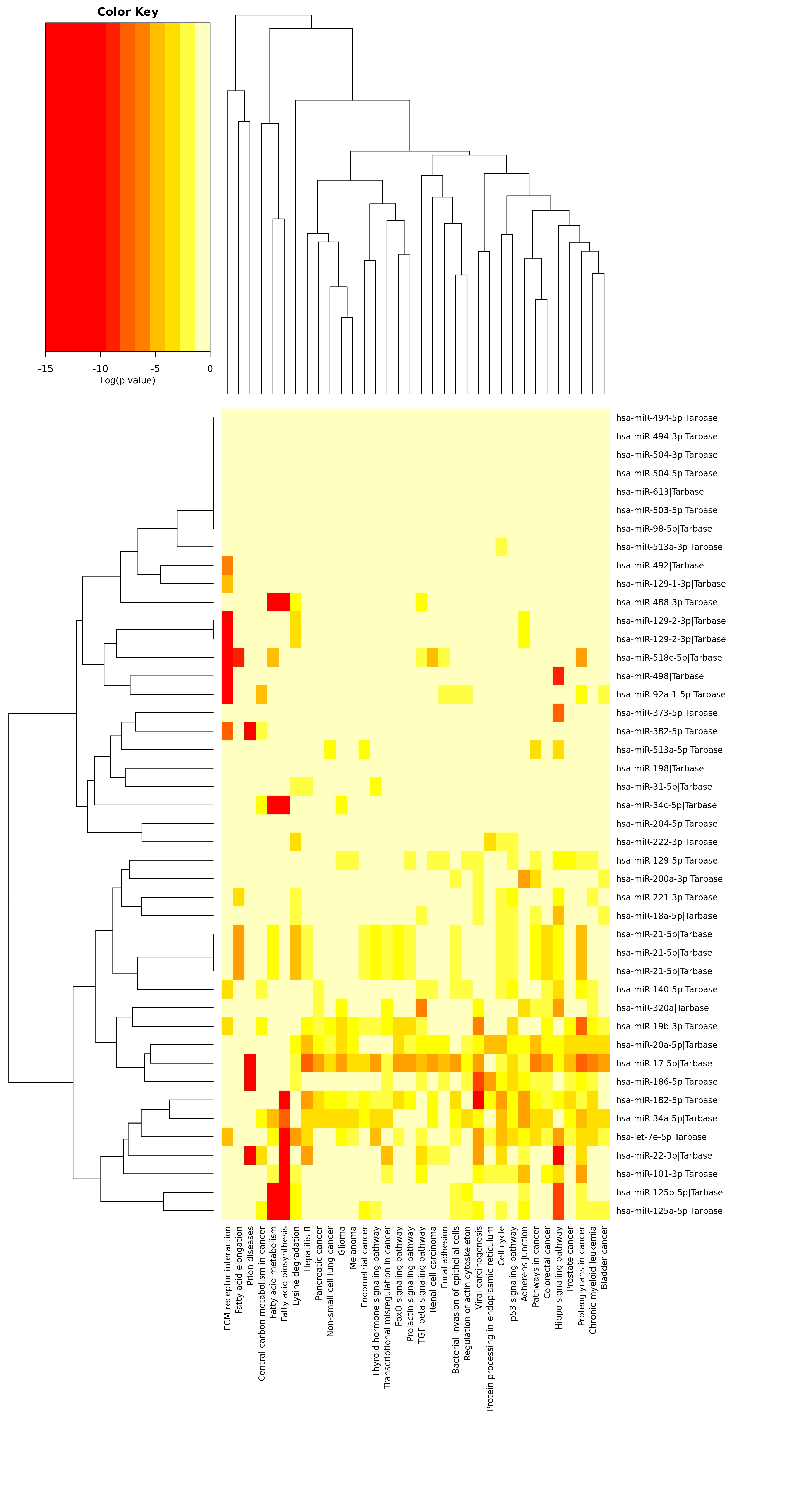
